# Mechanical stress and caveolin-1 control the release of extracellular vesicles with increased tumorigenic properties

**DOI:** 10.1101/2024.09.05.611225

**Authors:** Cristian Saquel, Louis Bochler, Ewan MacDonald, Celine Gracia, Christine Viaris de Lesegno, Frederik Verweij, Carlos Urena-Martin, Vanessa Masson, Damarys Loew, Graça Raposo, Jacky Goetz, Vincent Hyenne, Christophe Lamaze

## Abstract

Mechanical forces within the tumor microenvironment critically influence cancer progression, in part by modulating extracellular vesicle (EV) dynamics. However, the molecular mechanisms by which EVs respond to mechanical cues remain poorly understood. Here, we identify caveolin-1 (Cav1), a core structural component of caveolae, specialized membrane invaginations that act as mechanosensors, as a key regulator of EV release and composition under mechanical stress. We demonstrate that both 2D osmotic shock and 3D mechanical compression significantly enhance EV secretion from cancer cells in a Cav1-dependent manner. These EVs exhibit a distinct lipid profile and are selectively enriched in proteins involved in cell adhesion, motility, wound healing and extracellular matrix remodeling, cargo that is lost upon Cav1 deletion. Functionally, EVs secreted from mechanically stressed cells preferentially accumulate in the liver *in vivo* and promote breast cancer cell migration and invasion *in vitro*, effects abolished in the absence of Cav1. Together, these findings reveal a previously unrecognized role for Cav1 in coupling mechanical stress to EV biogenesis and cargo loading, thereby promoting metastatic traits. This study underscores a novel mechanism by which mechanical cues within tumor environment regulate EV-mediated communication and suggests that Cav1-enriched EVs may contribute to pre-metastatic niche formation.

## Introduction

Extracellular vesicles (EVs) are small membranous particles present in all bodily fluids, released into the extracellular space by all cell types studied thus far (van Niel et al., 2018; Welsh et al., 2024). In recent years, EVs have garnered significant attention due to their involvement in a wide range of physiological and pathological processes, including cancer (Yáñez-Mó et al., 2015; Becker et al., 2016). In the context of cancer, EVs contribute to various aspects of tumor progression and metastasis, such as promoting cell proliferation, invasion, angiogenesis, and immune evasion. Notably, it has been suggested that tumor cells release higher amounts of EVs compared to healthy cells (Logozzi et al., 2009). Tumor-derived EVs modulate the local tumor microenvironment and can also travel to distant organs, contributing to metastasis. Metastasis often exhibits organotropism, wherein cancer cells preferentially colonize specific organs due to molecular interactions and the characteristics of the local microenvironment. Similarly, EV-mediated communication in cancer, also display organotropism, as EVs preferentially target specific organs based on their molecular composition. This selective targeting promotes organ-specific metastatic progression by promoting the formation of pre-metastatic niches. This process, known as priming, involves the functional reprogramming of resident cells, to create a supportive environment for tumor cell colonization (Peinado et al., 2017; Ghoroghi et al., 2021a). EV-mediated organotropism is driven, in part, by the expression on their surface of molecules such as, integrins and adhesion proteins like CD146 (Ghoroghi et al., 2021b), which mediate the recruitment of EVs to specific tissues. These molecules likely interact with organ-specific extracellular matrix components or resident cell receptors, thereby determining EV tropism and guiding metastatic dissemination (Hoshino et al., 2015).

Caveolae are specialized, 60-80 nm cup-shaped invaginations of the plasma membrane found in most cell types (Lamaze et al., 2017, 2025). The structural and functional integrity of caveolae depends on both caveolins and cavins. Mutations or abnormal expression of caveolae components have been linked to various disorders, including cancer (Le Lay and Kurzchalia, 2005; Goetz et al., 2008; Singh and Lamaze, 2020). Notably, Cav1 has also been shown to regulate cholesterol levels in multivesicular bodies (MVBs), influencing membrane properties and cargo sorting into EVs (Albacete-Albacete et al., 2020). In 2011, a new function of caveolae was established as mechano-sensors and mechano-transducers, which are essential for cell mechano-protection (Sinha et al., 2011; Torrino et al., 2018; Dewulf et al., 2019; Kailasam Mani et al., 2025). As tumors progress and metastasize, their physical microenvironment undergoes significant changes, with evidence indicating that stromal caveolin-1 and caveolae play key roles in modulating tumor progression and metastatic dissemination (Goetz et al., 2011). Furthermore, tumor cells must continuously adapt and respond to these varying mechanical stresses encountered along the way (Follain et al., 2020; Gensbittel et al., 2021). Mechanical stress has also been shown to alter tumor extracellular vesicle secretion, though the underlying molecular mechanisms remain poorly understood (Guo et al., 2021; Wu et al., 2023; Sneider et al., 2024).

In this study, we explored the role of caveolae in EV dynamics in cancer cells subjected to mechanical strain using two distinct models; hypo-osmotic shock, which increases membrane tension through rapid cell swelling, and dextran-induced compression, applying external physical pressure onto cells. We observed a significant increase in the release of EVs under these two types of mechanical stress. The increase in EV production was strictly dependent on caveolin-1 (Cav1) and the endosomal sorting complex required for transport (ESCRT)-0 proteins. This specific subpopulation of Cav1-positive EVs exhibited a particular lipid and protein signature that was associated with increased liver tropism *in vivo* and the induction of metastatic traits in recipient cells, including increased migration and invasion *in vitro*. Our findings reveal a novel mechanism by which cancer cells exploit the mechanical properties of caveolae to adapt to the mechanical environment of tumors and promote the secretion of pro-metastatic EVs.

## Results

### Mechanical stress enhances EV release

To investigate the effect of mechanical stress on the release of EVs, we first applied a brief hypo-osmotic shock to increase membrane tension through cell swelling (**Figure S1A**). This biophysical perturbation mimics aspects of the mechanical stresses encountered by tumor cells and tissues *in vivo,* including those arising from interstitial fluid accumulation and elevated hydrostatic pressure within the tumor microenvironment (Northcott et al., 2018). The resulting increase in membrane tension leads to the flattening of caveolae and the release of caveolar components from the caveolar compartment (Sinha et al., 2011). EVs were purified from wild-type (WT) HeLa cells under resting or osmotic shock conditions using differential ultracentrifugation. Nanoparticle tracking analysis (NTA) of the size distribution revealed that most secreted EVs ranged between 50-200 nm (**Figure 1A**). Notably, cells subjected to hypo-osmotic shock exhibited a substantial 3.5-fold increase in secreted EVs (**Figure 1B**). To confirm that the observed effect was not specific to the two-dimensional hypo-osmotic shock model, we employed a three-dimensional mechanical compression model. This system utilized multicellular spheroids grown on an agarose bed in the presence of high molecular weight dextran (2 MDa), a sugar neutral to mammalian cell metabolism (Dolega et al., 2017) (**Figure S1B and S1C**). This 3D model recapitulates key features of the tumor microenvironment, where proliferating cells are subjected to increasing solid stress and confined growth due to limited extracellular space and compressive forces exerted by the surrounding stromal components. We found that this alternative type of mechanical strain also led to the disassembly of caveolae without affecting cell viability (**Figure S1D and S1E**). Dextran compression of spheroids induced a significant two-fold increase in the number of secreted EVs compared to resting spheroids (**Figure 1C**). To assess the purity and integrity of the released EVs, we used electron microscopy (EM). EM micrographs showed EVs with a smaller mean diameter (∼80 nm) compared to the average size measured by NTA (∼128 nm mean size), likely due to the difference in measuring techniques (**Figure 1D and 1E**). Western blot analysis confirmed the presence of *bonafide* EV markers, including CD63 and CD9 as previously described (Mathieu et al., 2019), although CD9 levels were consistently lower than those of CD63, suggesting a predominant endosomal origin rather than plasma membrane-derived EVs. The endoplasmic reticulum (ER) protein calnexin (CNX), commonly used as a negative marker, was present in cell extracts but absent in EVs, as expected. Additionally, the two main components of caveolae, Cav1 and cavin1, were detected in the EVs from WT HeLa cells (**Figure 1F**). Notably, Cav1 showed an approximate 2-fold enrichment in EVs from WT cells subjected to hypo-osmotic shock compared to those from resting cells (**Figure 1G**).

**Figure 1:**
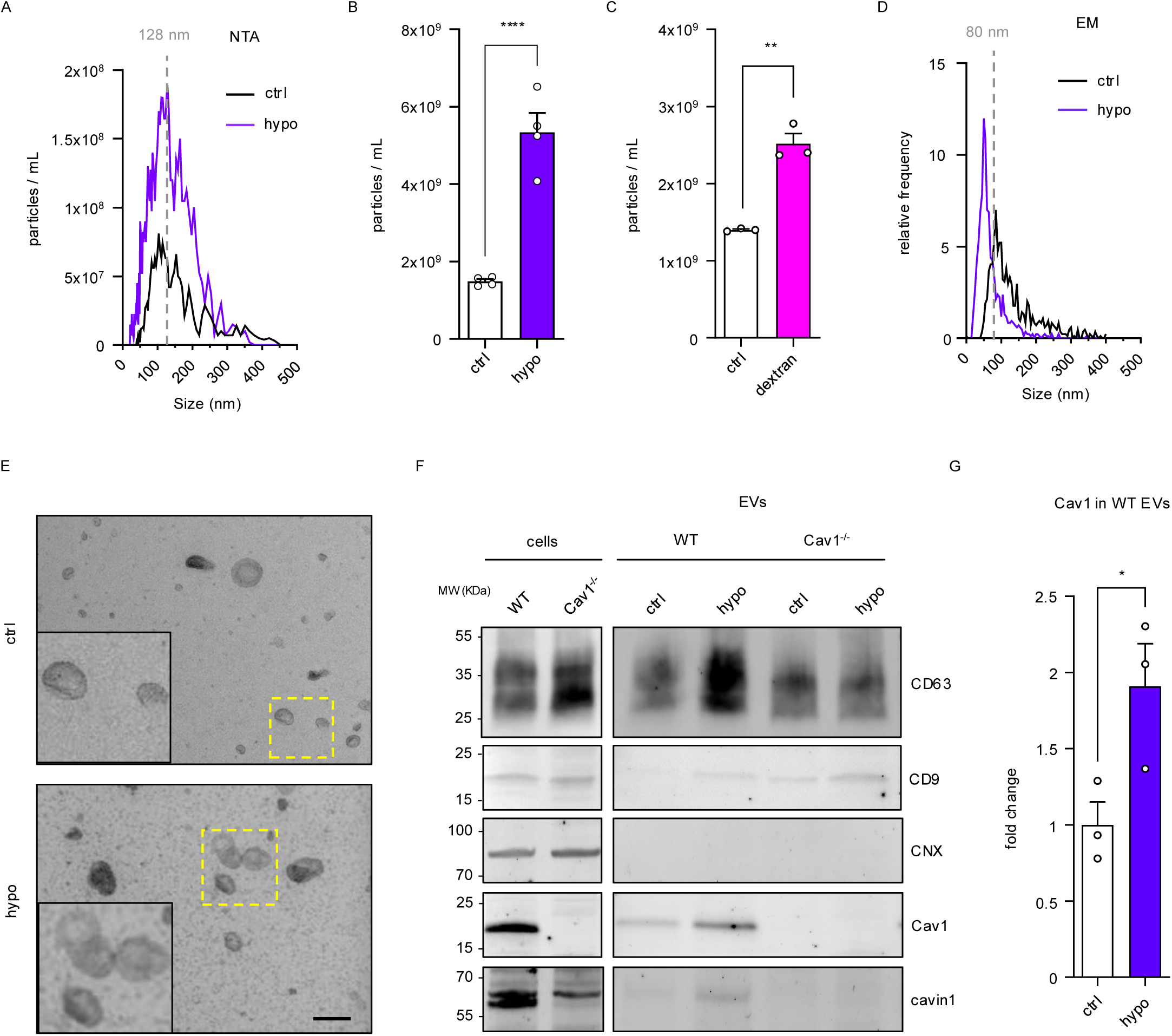
Mechanical stress increases EV secretion. **A.** Concentration (particles/ml) and size (nm) of EVs from WT HeLa cells in response to hypo-osmotic shock analyzed by NTA. **B.** Quantification of EVs released from WT HeLa cells in response to hypo-osmotic shock. Graph shows number of total secreted EVs as analyzed by NTA. **C.** Quantification of EVs released from WT HeLa cells in response to dextran compression. Graph shows number of total secreted EVs as analyzed by NTA. **D.** Vesicle concentration and size of EVs from WT HeLa cells in response to hypo-osmotic shock analyzed by EM. **E.** Representative EM images showing EVs isolated from WT HeLa cells in resting conditions or after hypo-osmotic shock. Scale bar 150 nm. **F.** Western Blot analysis of CD63, CD9, calnexin (CNX), Cav1 and cavin1 of WT or Cav1^-/-^ HeLa cell lysates and their corresponding purified EVs. **G.** Western Blot analysis of Cav1 in EVs from WT cells in resting or after mechanical stress conditions. Gels were loaded with proteins extracted from the same number of EVs. *: P < 0.05, ***: *P* < 0.01.

### Cav1 is required for mechanically induced EV release

Given the role of caveolae in mechano-sensing, we investigated their involvement in the increased release of EVs induced by mechanical stress. Remarkably, in HeLa cells stably knocked out for Cav1 expression, the hypo-osmotic no longer triggered an increase in EV secretion (**Figure 2A**). As with hypo-osmotic shock, the absence of Cav1 prevented this increase in EV secretion after mechanical compression (**Figure 2B**). Cavin1 is required for caveolae morphogenesis and stability at the plasma membrane. In the absence of cavin1, caveolae fail to form, and only Cav1 is present at the plasma membrane, albeit at lower expression levels (Drab et al., 2001; Hill et al., 2008). In HeLa cells stably knocked out for cavin1, mechanical stress still caused an increase in EV secretion, though this increase was significantly less pronounced than in WT HeLa cells (**Figure 2A and 2B**). Since cavin1 depletion is known to reduce Cav1 levels due to co-transcriptional regulation (Hill et al., 2008), we re-expressed Cav1 in Cavin1-deficient HeLa cells to restore its basal levels. Under these conditions, EV secretion returned to levels comparable to those induced by mechanical stress in WT HeLa cells (**Figure S2A**). Since cavin-1 deletion prevents caveolae assembly, these findings suggest that Cav1 itself, rather than budded caveolae, is required for the mechanically induced increase in EV secretion.

**Figure 2:**
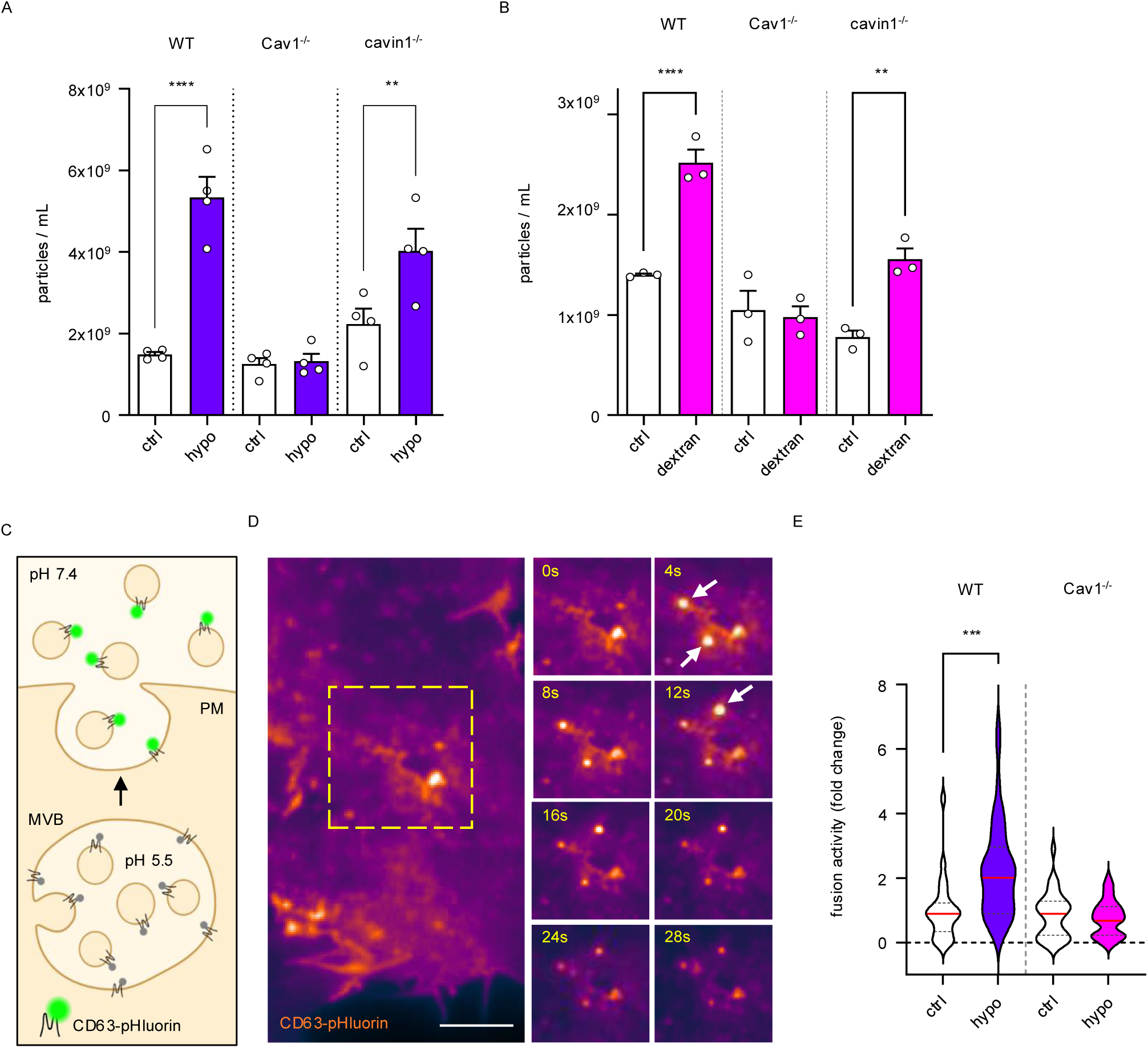
Cav1 is required for mechanically induced EV release. **A.** Quantification of EVs released by WT, Cav1^-/-^ and cavin1^-/-^ HeLa cells in response to hypo-osmotic shock. Graph shows the number of total secreted EVs quantified by NTA. **B.** Quantification of EVs released by WT, Cav1^-/-^ and cavin1^-/-^ HeLa cells in response to dextran compression. Graph shows the number of total secreted EVs quantified by NTA. **C.** Schematic representation of the reporter CD63-pHluorin for MVB-PM fusion visualization. **D.** Representative time-lapse imaging of an MVB-PM fusion event and release of exosomes, indicating fluorescence signal duration characteristic of this type of fusion event. Arrows indicate fusion events. **F.** Quantification of MVB fusion activity (events per cell and time) of WT or Cav1^-/-^ HeLa cells in response to hypo-osmotic shock. **** *P <* 0.0001, *** *P <* 0.001, ** *P* < 0.01.

We next investigated whether this process is conserved in triple negative breast cancer (TNBC) cells. To this end, we examined two TNBC cell lines, Hs578t and MDA-MB-231, both characterized by aggressive phenotypes. In both cell lines, hypo-osmotic shock and dextran-induced compression significantly increased EV release, as measured by NTA. Notably, this mechanically increased EV secretion was abolished in Cav1-deficient cells and restored upon Cav1 re-expression, highlighting a conserved role for Cav1 in mechanotransduction-driven EV release across distinct cancer cell types (**Figure S2B-D**). Different mechanisms may contribute to increased EV production, including enhanced secretion by producing cells or impaired reuptake by recipient cells. To address the former possibility, we specifically measured EV release using a CD63-pHluorin reporter-construct as previously described (Verweij et al., 2018). CD63-pHluorin is a pH-sensitive reporter protein that fluoresces once the pH inside endosomal acidic compartments is neutralized, such as the pH ∼5.5 of multivesicular bodies (MVB), containing intraluminal vesicles (ILV), the precursors of exosomes (**Figure 2C**). This optical sensor thus allows visualization of MVB fusion at the plasma membrane and the subsequent release of ILVs as exosomes into the pH-neutral extracellular medium from live cells. In line with EV secretion results measured by NTA, endosome-dependent exosomal release activity in HeLa cells increased following osmotic shock (**Figure 2D and 2E**). This increase in fusion activity after mechanical stress was absent in HeLa Cav1^-/-^ cells, confirming that the enhanced EV detection was due to increased EV release by the producing cells. Altogether, these findings reveal a novel Cav1-dependent mechanism by which hypo-osmotic shock and 3D mechanical compression dramatically increase the release of small EVs (i.e. exosomes) from cells.

### Mechanical EV release requires the endosomal ESCRT-0 complex and Cav1 ubiquitination

We next explored the mechanism by which the MVB pathway is influenced by mechanical stress or Cav1 depletion. First, we used EM to analyze the characteristics of MVBs in Hs578t cell spheroids under resting conditions, as well as after 5 min, 1 day, and 5 days of dextran compression (**Figure 3A**). Although the diameter of MVBs after mechanical compression did not show significant changes over time (**Figure 3B**), the overall number of MVB-like organelles in compressed Hs578t spheroids was significantly reduced compared to resting spheroids across all compression durations (**Figure 3C**). Additionally, in HeLa cells subjected to osmotic shock, we observed enlarged CD63-positive MVBs decorated with Cav1 (**Figure S1F and S1G**). This result was confirmed by EM in MLEC cells that are particularly abundant in caveolae since a hypo-osmotic shock led to an increase in Cav1-immunogold labelling in MVB structures (**Figure S1H**).

**Figure 3:**
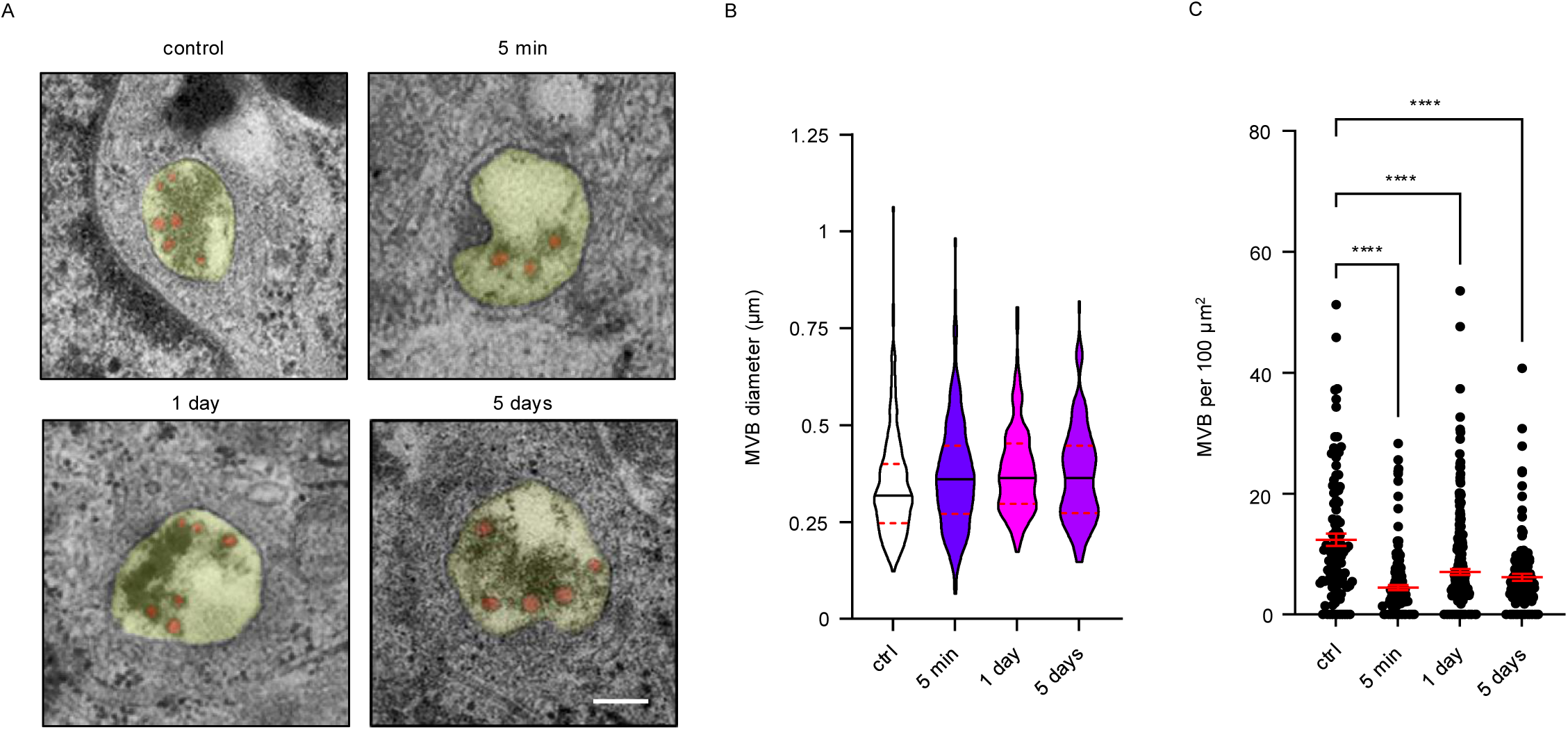
Mechanical stress reduces the amount of multivesicular bodies. **A.** Representative EM image of MVBs in an Hs578t cell spheroid in control state, 5 min, 1 day and 5 days after dextran compression. Scale bar 50 nm. **B.** Quantification of MVB diameter in Hs578t cell spheroids after 5 min, 1 day or 5 days of dextran compression. **C.** Quantification of the number of MVBs in Hs578t cell spheroids after 5 min, 1 day or 5 days of dextran compression, represented by area. **** *P* < 0.0001.

We next assessed the protein levels of ESCRT pathway components, a key machinery involved in MVB biogenesis and exosome production. The signal-transducing adaptor molecule (STAM) binds to hepatocyte growth factor-regulated tyrosine kinase substrate (HRS) to form the ESCRT-0 complex at early endosomes (Migliano et al., 2022). Using western blot analysis, we found that both HRS and STAM were significantly downregulated, with their expression reduced to approximately 45% in the absence of Cav1 (**Figure 4A and 4B**). Next, we tested the effect of ESCRT-0 downregulation on the mechanically induced increase in EV release. Treating WT HeLa cells with siRNA targeting HRS led to efficient downregulation of HRS (>90%) and a milder downregulation of STAM (∼50%) (**Figure 4C**). We then purified EVs from HRS-downregulated cells subjected to hypo-osmotic shock. NTA revealed a significant reduction in secreted EVs in HeLa cells treated with siRNA against HRS (**Figure 4D**), in line with previous reports implicating HRS in ILV biogenesis (Colombo et al., 2013, 2014). Notably, mechanical stress failed to increase the number of secreted EVs in HRS-downregulated HeLa cells, indicating that the mechanically induced EV release also depends on ESCRT-0. This suggests a co-regulation of this effect by both Cav1 and ESCRT-0 (**Figure 4D**).

**Figure 4:**
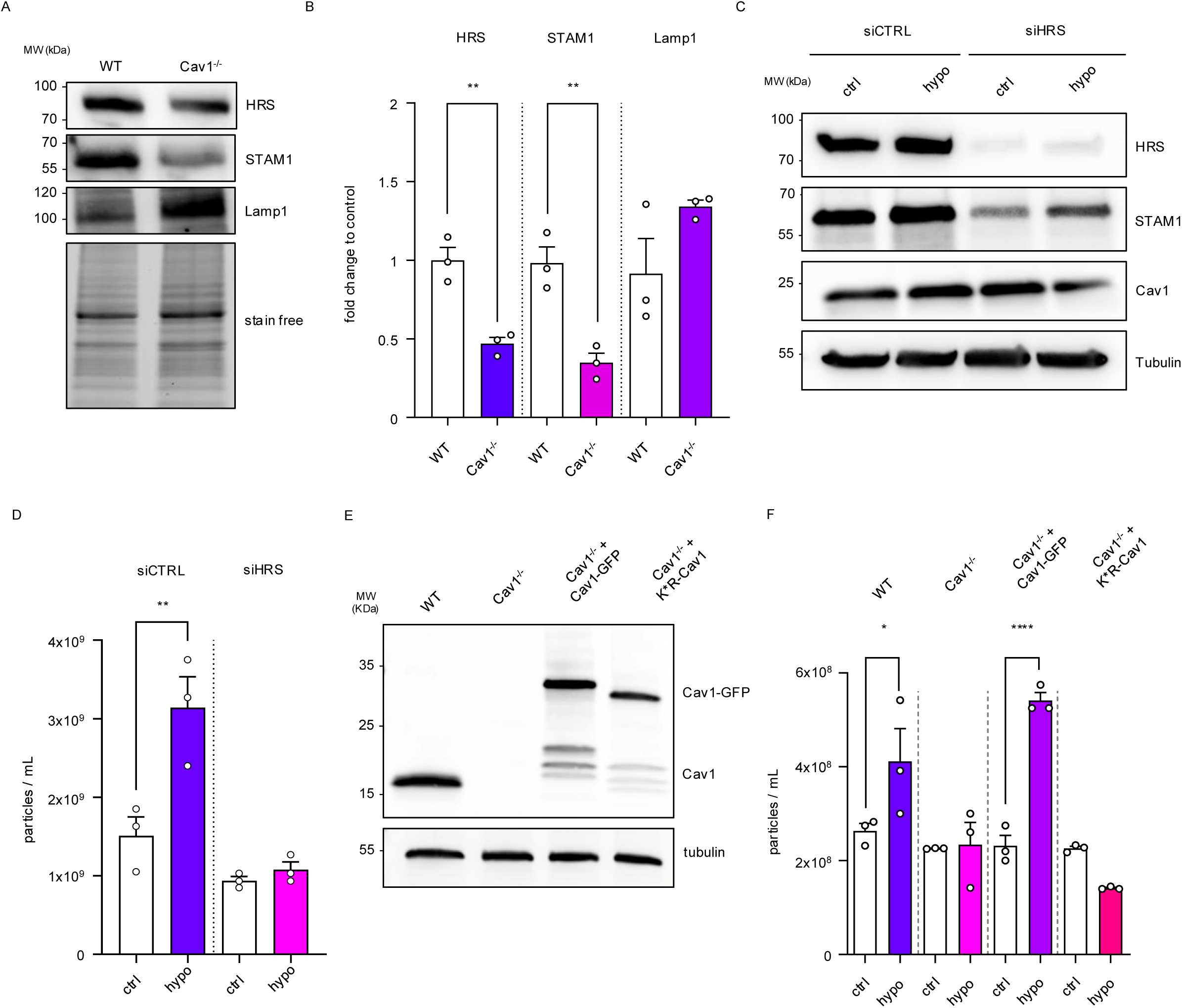
ESCRT-0 and Cav1 ubiquitination are required for mechanically induced EV release. **A.** Western Blot analysis of HRS, STAM, and Lamp1 in WT or Cav1^-/-^ HeLa cells lysates. **B.** Protein signal quantification graphs normalized to the stain free gel. **C.** Cell lysate Western Blot analysis of HRS, STAM, Cav1 and tubulin in HeLa cells treated with siRNA control or siRNA against HRS. **D.** Quantification of EVs released by HeLa cells treated with siRNA control or siRNA against HRS in resting conditions or after mechanical stress. Graph shows the number of total secreted EVs quantified by NTA. **E.** Western Blot analysis of Cav1 and tubulin in WT, Cav1^-/-^, Cav1^-/-^ transfected with Cav1-GFP or Cav1^-/-^ transfected with Cav1-K*R HeLa cells. **F.** Quantification of EVs released by WT, Cav1^-/-^, Cav1^-/-^ transfected with Cav1-GFP or Cav1^-/-^ transfected with Cav1-K*R HeLa cells. Graph shows the number of total secreted EVs quantified by NTA. **** *P* < 0.0001, ** *P* < 0.01, * *P* < 0.05.

In the ESCRT-0 complex, both HRS and STAM, contain ubiquitin binding sites that promote the recruitment of EV cargo to the early endosome by recognizing ubiquitination in target proteins (Bache et al., 2003). To investigate the role of Cav1 ubiquitination in the Cav1-mediated enhancement of EV release after mechanical stress, we expressed in Cav1^-/-^ HeLa cells a ubiquitination resistant mutant of Cav1 (K*R-Cav1), in which lysines were replaced by arginines, which prevented Cav1 ubiquitination (**Figure 4E**). Our results show that expression of this ubiquitin-resistant Cav1 significantly hindered EV release from HeLa cells upon mechanical stress, as compared to Cav1^-/-^ HeLa cells expressing a functional GFP-tagged Cav1 (GFP-Cav1) serving as control. This remarkable reduction underlines the importance of Cav1 ubiquitination in promoting the stimulated release of EVs of HeLa cells under mechanical stress conditions (**Figure 4F**).

### Mechanical stress drives the release of EVs with a distinct lipid and protein signature

Caveolae organize lipid nanodomains at the plasma membrane that are enriched in cholesterol and sphingolipids, and they play a central role in regulating lipid homeostasis (Sonnino and Prinetti, 2009; Krishna and Sengupta, 2019; Prakash et al., 2021). We investigated whether mechanical stress and Cav1 depletion affect the lipid composition of EVs. To this end, we performed lipidomic analysis of EVs purified from WT and Cav1^-/-^ HeLa cells under resting conditions and after exposure to mechanical stress. Mass spectrometry (MS) revealed significant differences in the lipid composition of EVs between WT and Cav1-depleted HeLa cells, as well as between EVs derived from resting and mechanically stressed HeLa cells.

Mechanical stress specifically enriched certain lipid species in EVs, including phosphatidylcholine, O-phosphatidylcholine, phosphatidylethanolamine, O-phosphatidylethanolamine, and phosphatidylinositol. In contrast, mechanical stress resulted in a decrease in cholesterol and ceramide content in EVs. Cav1 depletion in cells led to an increase of lactosylceramide (LacCer), alongside a decrease in ceramide, the globoside Gb4 and the ganglioside GM2 in EVs (**Figure 5A-C**).

**Figure 5:**
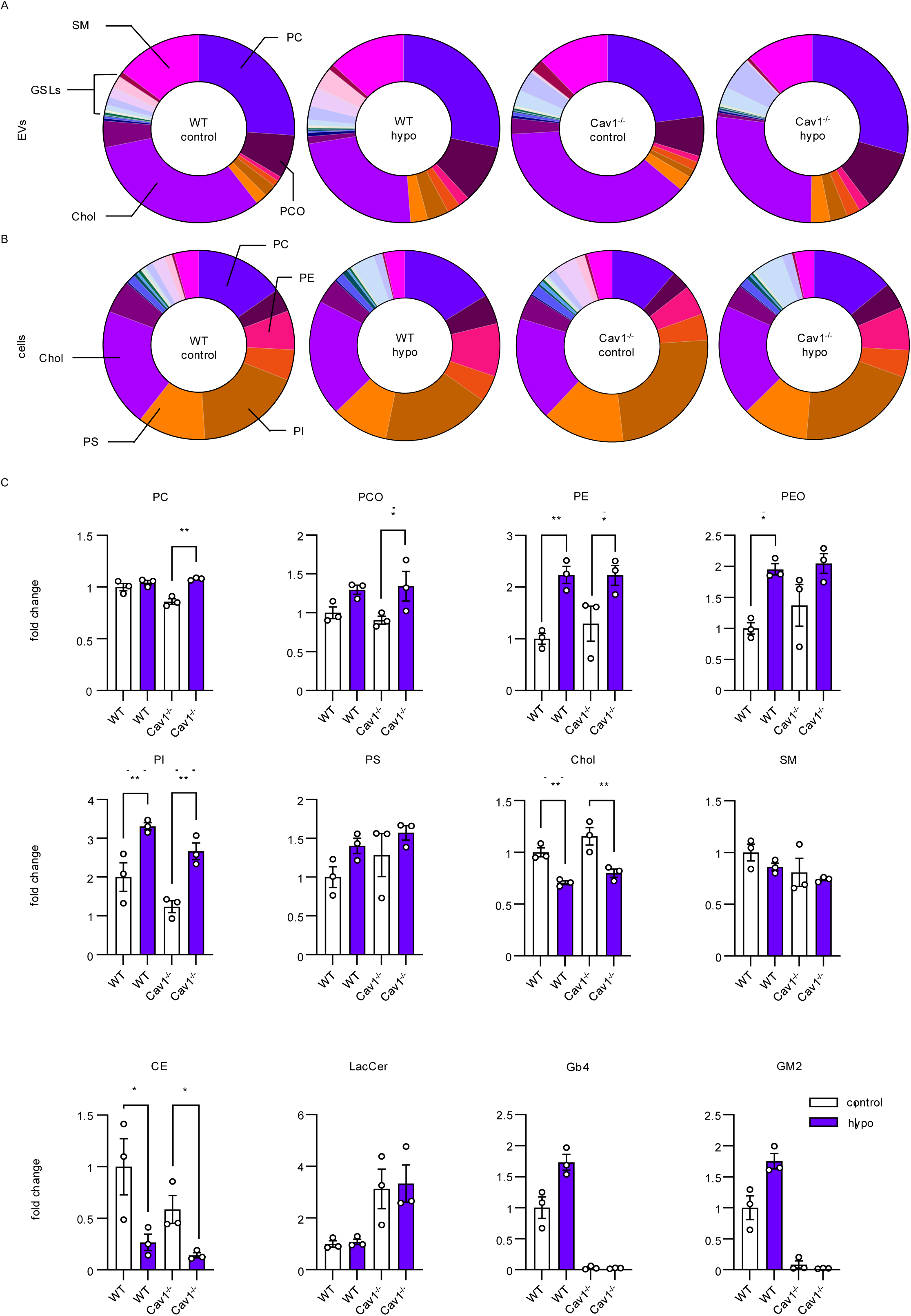
Mechanical stress and Cav1 modulate the lipid composition of EVs. **A.** Mol percentile of each lipid class in EVs from WT or Cav1^-/-^ HeLa cells in resting conditions or after mechanical stress. **B.** Mol percentile of each lipid class in lysates from WT or Cav1^-/-^ HeLa cells in resting conditions or after mechanical stress. **C.** Comparison of the fold change in lipid classes between EVs from WT or Cav1^-/-^ HeLa cells in resting conditions or after mechanical stress. ** *P* < 0.01, * *P* < 0.05.

Cav1 has been implicated in the sorting of various proteins into EVs. Recent studies have demonstrated that Cav1-dependent sorting of specific ECM and adhesion molecules, such as tenascin-C, fibronectin, and CD146, into EVs can promote malignancy in recipient cells (Albacete-Albacete et al., 2020; Campos et al., 2018, 2023; Ghoroghi et al., 2021b). To assess whether mechanical stress alters the protein content of EVs, particularly proteins involved in cell adhesion and ECM organization, we performed MS-based proteomic analysis of EVs from WT and Cav1-depleted cells under both mechanical stress and resting conditions. Our analysis revealed notable differences between EVs from cells in resting conditions and mechanically stressed, as well as between WT and Cav1-depleted cells. Qualitative analysis, focusing on proteins quantified with at least two distinct peptides, showed that while most proteins were shared across all four conditions, each condition also contained a distinct subset of unique proteins (**Figure 6A**). Principal component analysis (PCA) revealed a clear separation between EVs from WT cells and Cav1-depleted cells. Interestingly, EVs from control and mechanically stressed Cav1-depleted cells clustered closer together (**Figure 6B**).

**Figure 6:**
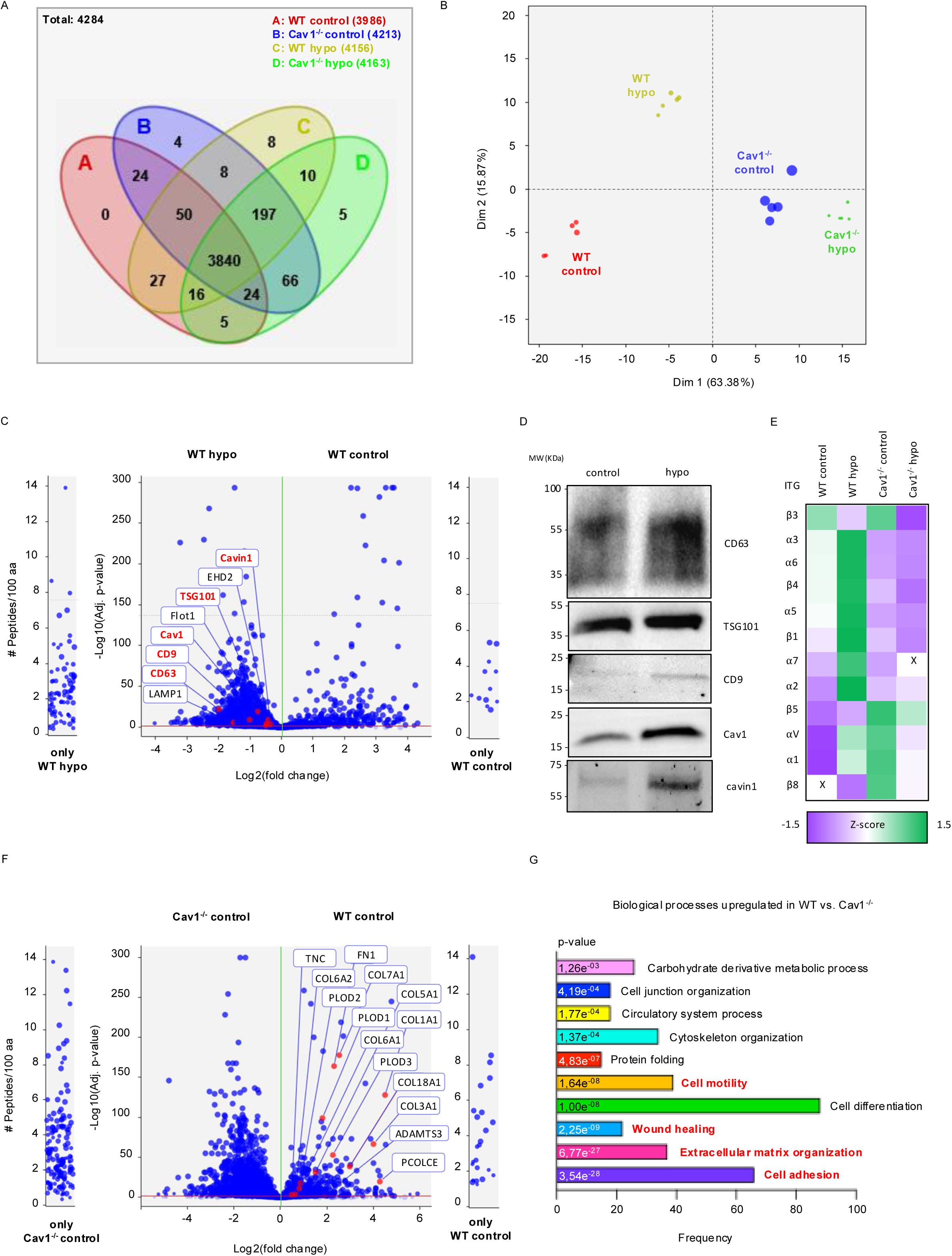
Mechanical stress and Cav1 modulate the protein cargo of EVs involved in adhesion and ECM remodeling. **A.** Venn diagram illustrating the qualitative distribution of the proteins quantified in each condition by at least two distinct peptides. **B.** PCA analysis showing the quantitative comparison and separation of fractions. **C.** Volcano plot displaying the quantitative analysis of proteins in WT control compared to WT hypo HeLa EVs. The horizontal red line indicates an adjusted *P* value of 0.05, and the vertical green line indicates an absolute fold-change of 1. Proteins highlighted are selected as potential specific markers of interest. **D.** Western blot validation of the highlighted proteins of interest from panel C. **E.** Heatmap displaying the Z-score of integrins in WT control, WT hypo, Cav1^-/-^ control and Cav1^-/-^ hypo HeLa EVs. **F.** Volcano plot showing the quantitative analysis of proteins in WT control compared to Cav1^-/-^. The horizontal red line indicates an adjusted *P* value of 0.05, and the vertical green line indicates an absolute fold-change of 1. Highlighted proteins are selected as potential specific markers of interest. **G.** Quantitative Gene Ontology (GO) term enrichment analysis for WT control versus Cav1^-/-^ control fractions. The top 10 significantly enriched GO categories under Biological Processes are indicated.

A volcano plot representing the relative abundance of proteins in EVs from WT cells under control or mechanical stress conditions revealed significant enrichment (*P* < 0.05) of several proteins in EVs released after mechanical stress. Notably, these EVs were enriched in classic EV markers such as CD63, CD9, TSG101, Lamp1 and Flotillin-1. In addition, there was a marked enrichment of caveolar proteins, including Cav1, cavin1, and EHD2 (**Figure 6C**). The presence of several of these EV and caveolar proteins was further validated by western blot analysis (**Figure 6D**). Interestingly, we observed substantial differences in integrin enrichment between EV samples. Most integrins were significantly more abundant in EVs from WT cells after osmotic shock compared to those from WT cells in resting conditions (**Figure 6E**). Furthermore, a comparison of EVs from WT and Cav1-depleted cells under control conditions showed significant decrease (*P* < 0.05) of various proteins associated with ECM remodeling and ECM components, such as collagenases, metalloproteinases, tenascin-C, fibronectin, collagen, and laminin, among others (**Figure 6F**).

To further analyze the potential signaling pathways down-regulated in EVs from WT or Cav1-depleted cells, we performed gene ontology (GO) enrichment analysis using the myProMS software (https://github.com/bioinfo-pf-curie/myproms). Proteins were classified by biological processes. In agreement with the specific proteins enriched in EVs from WT cells, the analysis revealed that biological processes such as cell adhesion, ECM organization, wound healing, and cell motility were significantly enhanced in EVs from WT cells compared to those from Cav1-depleted cells (**Figure 6G**).

### Mechanical stress promotes liver tropism of EVs via Cav1

EV organotropism is shaped by various molecules present on their surface, including integrins, which contribute to direct interactions with target tissues. Prior studies have demonstrated that EV-associated integrins such as αVβ5 confer liver tropism, whereas α6β4 and α6β1 promote localizations to the lungs (Hoshino et al., 2015). Our MS-based proteomic analysis of HeLa-derived EVs revealed distinct integrin signatures, which we validated in EVs from Hs578t cells by western blot, revealing increased expression of αV and β5 integrins upon mechanical stress (**Figure S3A**). These findings led us to investigate whether mechanical stress and Cav1 expression cooperatively modulate EV biodistribution *in vivo*.

To test this, we intravenously injected equal amounts of EVs derived from WT or Cav1^-/-^ Hs578t cells, isolated under resting or mechanically stressed conditions, into mice. After 45 minutes, major organs were harvested and imaged to assess EV biodistribution (**Figure 7A**).

**Figure 7:**
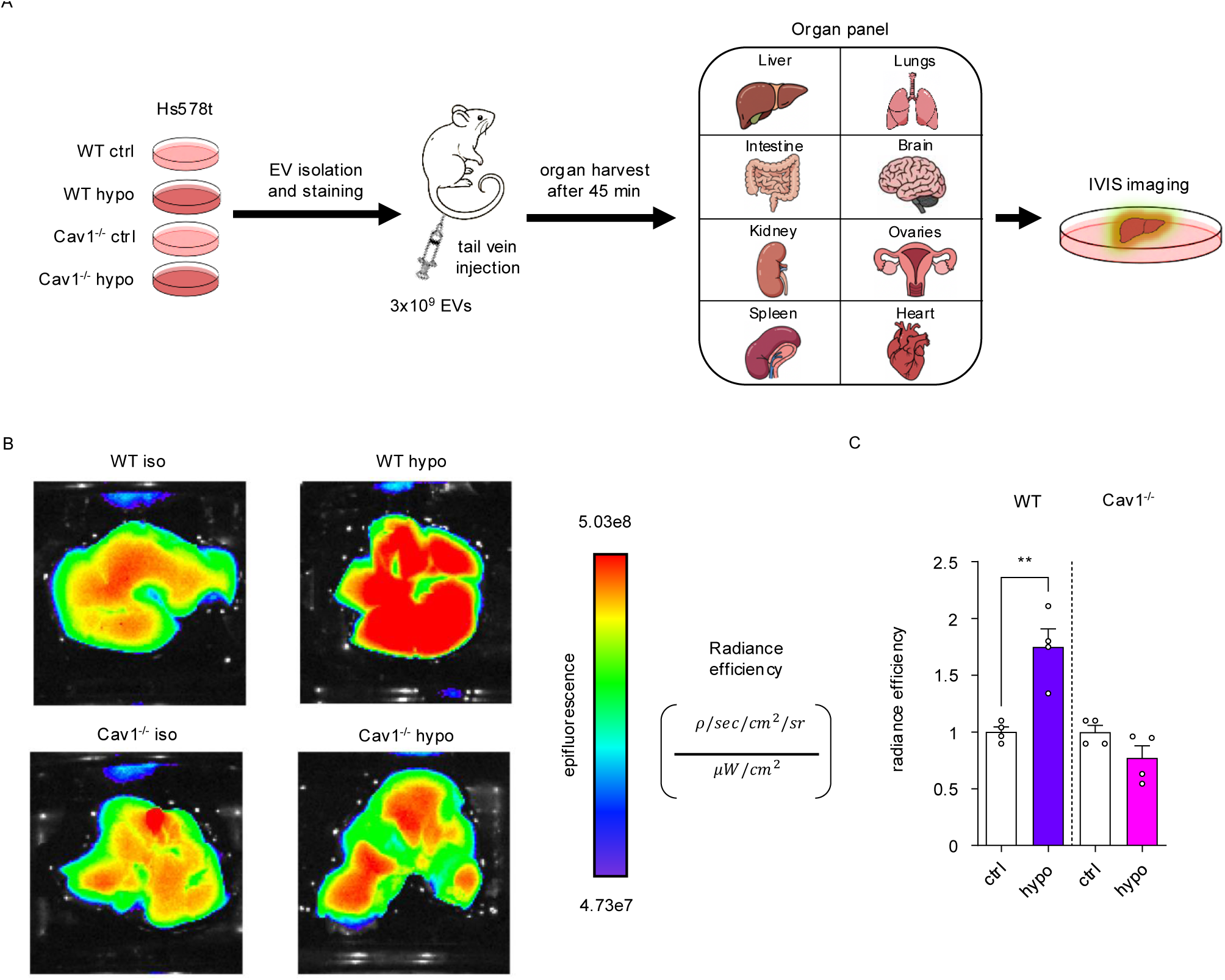
Mechanical stress enhances liver tropism of EVs in a Cav1-dependent manner. **A.** Schematic representation of the experimental workflow. EVs were isolated from WT and Cav1^-/-^ HS578t breast cancer cells under resting or mechanically stressed conditions. 3x10^9^ EVs were fluorescently labeled and injected into the tail vein of mice. After 45 minutes, liver, lungs, intestine, brain, kidneys, ovaries, spleen and heart were harvested, and IVIS imaging was performed to assess EV biodistribution. Fluorescence intensity indicates accumulation of EVs in the corresponding organ. **B.** Representative epifluorescence images of the liver showing EV accumulation. **C.** Quantification of radiance efficiency. Results were normalized in both WT and Cav1^-/-^ conditions to their respective control in resting conditions. ** *P* < 0.01.

EVs from mechanically stressed WT cells exhibited significantly increased accumulation in the liver compared to all other conditions. This effect was markedly reduced in the absence of Cav1, with liver fluorescence levels from Cav1^-/-^ mechanically stressed EVs comparable to control EVs (**Figure 7B and 7C**). In contrast, no significant differences in EV distribution were observed in other organs, including the spleen, heart, lungs, intestines, brain, kidneys, and ovaries, highlighting the specificity of liver accumulation in the WT mechanically stressed EV group (**Figure S3B-I**).

These results suggest that mechanical stress induces Cav1-dependent alterations in EV composition that enhance liver tropism, potentially through modulation of integrin content. Cav1 deficiency abrogates this effect, underscoring its essential role in the mechanotransduction-driven modulation of EV biodistribution.

### EVs from mechanically stressed cells promote cancer cell migration and invasion

The observed changes in the lipid composition and protein cargo of EVs suggests that these vesicles may deliver bioactive cues capable of modulating the behavior of recipient cells. Metastatic progression depends, at least in part, by the ability of tumor cells to migrate and invade surrounding tissues, we next investigated whether EVs produced under mechanical stress could directly promote these pro-metastatic traits. To this end, we investigated the functional consequences of EV uptake in recipient cells using cell proliferation and invasion assays. Proliferation was assessed using a wound-healing assay (Ilina and Friedl, 2009), where we found that EVs from mechanically stressed cells accelerated wound closure in WT Hs578t cells compared to EVs from resting cells (**Figure 8A-C**). Importantly, this effect was dependent on Cav1, as EVs derived from Cav1-depleted cells did not promote wound closure under either condition. To evaluate the invasive potential of these EVs, Hs578T cells were pre-incubated with EVs for 24 h prior to seeding on a thin layer of Matrigel, which mimics the ECM and contains collagen type IV, entactin, laminin, and heparan sulfate proteoglycan. EVs from mechanically stressed WT cells significantly enhanced the invasion of Hs578t cells through the ECM, while EVs from resting cells induced a slightly more modest increase. In contrast, EVs from Cav1-deleted cells failed to promote invasion, underscoring the requirement of Cav1 for this functional effect (**Figure 8D**).

**Figure 8:**
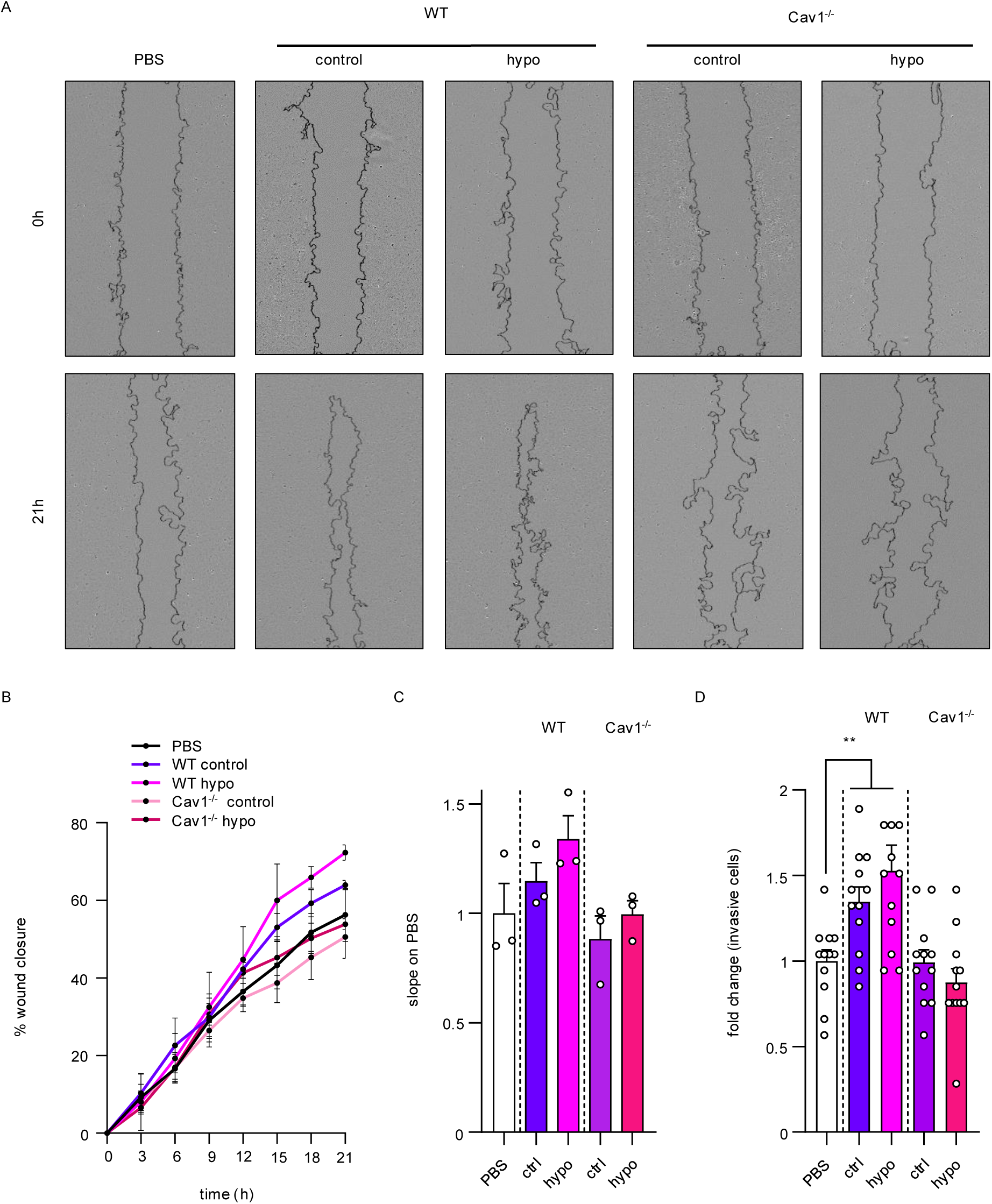
EVs from mechanically stressed cells promote migration and invasion in a Cav1-dependent manner. **A.** Representative images of wound healing assays on Hs578t cells pre-treated with the same quantity of Hs578t EVs from WT or Cav1^-/-^ cells in resting conditions or after exposure to mechanical stress at times 0 and 21 hours. **B.** Wound closure quantification represented in percentage over time. **C.** Quantification of the slope of the wound closure curves. **D.** Quantification of invasive Hs578t cells pre-treated with the same quantity of Hs578t EVs from WT or Cav1^-/-^ cells in resting conditions or after exposure to mechanical stress. ** *P* < 0.01.

To determine whether differences in EV uptake underlie variations in recipient cell migration and invasion, equal numbers of EVs from WT and Cav1-depleted HeLa cells were labeled with CFDA-SE, a membrane-permeable dye that fluoresces upon esterase cleavage inside EVs to form CFSE (Morales-Kastresana et al., 2017). After a 4-hour incubation with Hs578t cells, both EV populations showed comparable uptake levels (**Figure S4A and S4B**), indicating that functional differences arise from intrinsic EV properties rather than uptake efficiency.

In summary, our data indicate that EVs secreted by Hs578t cells in response to mechanical stress enhance the migratory and invasive capabilities of Hs578t cells through the ECM in a Cav1-dependent manner, thereby increasing their tumorigenic potential. EVs from resting Hs578t cells present a slightly reduced migratory and invasive inducing capabilities over Hs578t cells.

## Discussion

In this study, we identified Cav1, the principal structural constituent of caveolae, as a central regulator of extracellular vesicle biogenesis in response to mechanical stress. EVs secreted under these conditions exhibit distinct lipid and protein signatures, demonstrate a tropism for the liver *in vivo,* and enhance migration and invasion *in vitro*—effects that are critically dependent on Cav1.

EVs isolated from HeLa WT, Cav1^-/-^, and cavin1^-/-^ cells ranged up to approximately 350 nm in size, with over 80% measuring below 200 nm. Their exosomal nature was confirmed by monitoring MVB-plasma membrane fusion using a pHluorin-CD63 reporter and demonstrating dependence on a functional ESCRT-0 complex. In addition to canonical EV markers such as CD63 and CD9 (Keerthikumar et al., 2016; Mathieu et al., 2019), Cav1 and cavin1 were detected in EVs from WT HeLa cells, in line with previous reports (Campos et al., 2018; Logozzi et al., 2009).

The molecular mechanisms by which mechanical stress enhances EV secretion through Cav1 remain incompletely understood. Our data indicate that both Cav1 and ESCRT-0 are required for this response. Previous studies have implicated interactions between Cav1 and ESCRT machinery, including HRS-dependent targeting of Cav1 to intraluminal vesicles (ILVs) within MVBs (Hayer et al., 2010). Cav1 overexpression also promotes its accumulation in endosomal compartments, suggesting a role in EV biogenesis.

Given caveolae’s established role in mechano-sensing and membrane protection, we hypothesized that Cav1 levels modulate cancer cell response to mechanical stress (Singh and Lamaze, 2020). This is particularly relevant during tumor progression, when increasing physical forces accompany tissue stiffening and compression (Northcott et al., 2018). Cav1 is known to promote metastatic-related traits, such as cell migration, invasion and ECM stiffening (Goetz et al., 2011; Urra et al., 2012; Díaz et al., 2014), though the mechanisms remain unclear. The presence of Cav1 in EVs from breast cancer cells was associated with increased migration and invasion of breast cancer recipient cells (Campos et al., 2018). Our data show that EVs derived from mechanically stressed HeLa cells enhanced migration and invasion of Hs578t breast cancer cells. This effect was further enhanced with incubation with EVs secreted under mechanical stress. This effect was dependent on the presence of Cav1 in donor cells, suggesting that Cav1 is necessary for generating a metastasis-promoting EV subpopulation. Importantly, because Hs578t cells endogenously express Cav1, these effects result not solely from Cav1 transfer but from Cav1-dependent modifications in EV composition induced by mechanical stress.

To investigate the nature of these changes, we considered Cav1’s role in lipid clustering and nanodomain organization at the plasma membrane (Pohl et al., 2002; Pol et al., 2005; Doktorova et al., 2025), as well as in protein sorting into EVs (Campos et al., 2018, 2023). Moreover, Cav1 has been shown to act as a cholesterol rheostat in MVBs, regulating cholesterol content and thereby influencing membrane properties and cargo sorting (Albacete-Albacete et al., 2020). Mechanical forces are known to induce lipid phase separation in giant unilamellar vesicles (GUV) (Colom et al., 2018). Lipidomic profiling revealed significant differences in EV lipid composition between WT and Cav1-depleted cells and between resting and mechanically stressed conditions. Notably, mechanically stressed WT cells produced EVs enriched in globoside Gb4 and ganglioside GM2, two glycosphingolipids implicated in cancer progression Gb4 activates Raf-MEK-ERK signaling to drive proliferation (Park et al., 2012), while GM2 is overexpressed in tumors and promotes oncogenic signaling (Sasaki et al., 2021). These findings support a model in which mechanical stress promotes lipid reorganization at the MVB membrane, influencing both lipid and protein cargo selection.

MS-based proteomic analysis further revealed mechanical stress- and Cav1-dependent enrichment of proteins involved in ECM remodeling and cell adhesion including tenascin-C and fibronectin, which have been implicated in metastatic niche formation (Dourado et al., 2019; Campos et al., 2023). Additionally, mechanical stress altered the EV content of integrins, which are known to mediate organotropism and uptake by recipient cells (Hoshino et al., 2015). These alterations likely contribute to the liver-tropic behavior of EVs observed *in vivo*.

Mechanistically, we link increased EV production to the dynamic response of caveolae under mechanical stress. We and others have demonstrated that caveolae flatten and disassemble to buffer membrane tension and prevent rupture (Sinha et al., 2011), releasing caveolae coat components into the cytoplasm and non-caveolar Cav1 at the plasma membrane (Torrino et al., 2018; Kailasam Mani et al., 2025). Our data indicate that this pool of Cav1 is subsequently internalized, trafficked to MVBs, and incorporated into ILVs, culminating in secretion via EVs. Cav1-deficient cells fail to mount this response, while cavin1-deficient cells, which have only non-caveolar Cav1, can still secrete EVs, albeit at reduced levels. These data underscore a correlation between mechanical stress-induced EV production and Cav1 availability in the endosomal network. Moreover, we found that Cav1 ubiquitination is required for the increase in EV release observed under mechanical stress. Inhibition of this post-translational modification abrogated stress-induced EV secretion, highlighting its essential role in regulating the biogenesis of EVs in a Cav1-dependent manner.

Importantly, EVs from mechanically stressed breast cancer cells preferentially accumulated in the liver, a phenomenon likely driven by mechanical stress-induced changes in EV composition, particularly alterations in integrin expression and adhesion proteins, which are known to dictate organ-specific EV uptake (Hoshino et al., 2015; Ghoroghi et al., 2021; Singh et al., 2024). Our data revealed a specific enrichment of integrins αV and β5 in EVs in mechanically-stressed WT cells—two integrins previously associated with liver tropism (Hoshino et al., 2015). This distinct integrin profile, along with Cav1-regulated enrichment of other adhesion molecules, likely contributes to the selective uptake of these EVs by hepatic cells. Notably, the absence of Cav1 abolished this effect, reinforcing its role in sorting key adhesion molecules that determine EV biodistribution. The liver is a frequent site of metastasis in aggressive cancers, and recent work has revealed that tumor-derived EVs are able to induce hepatic reprogramming, generating a pro-inflammatory environment, dysregulating liver metabolism, and promoting fatty liver formation (Wang et al., 2023). Our data suggest that Cav1-enriched EVs released under mechanical stress may contribute to the establishment of pre-metastatic niches in this organ. Given the increasing mechanical rigidity of the tumor microenvironment over time (Northcott et al., 2018; Gensbittel et al., 2021), this mechanism could be further amplified during cancer progression. Consistent with this notion, recent studies have shown that extracellular matrix stiffening not only enhances EV secretion but also alters their content, leading to promoting tumor growth and metastasis (Wu et al., 2023; Sneider et al., 2024).

We validated the functional impact of these EVs by demonstrating that mechanically stressed Hs578T WT released EVs that significantly enhanced migration and invasion *in vitro*, compared to EVs from resting or Cav1-deficient cells. These results underscore the ability of mechanical cues to modulate EV-mediated intercellular communication in a manner that promotes malignancy.

In conclusion, our study identifies Cav1 as a critical regulator of EV biogenesis and cargo sorting in response to mechanical stress, linking biomechanical forces to enhanced tumorigenic potential via EV-mediated communication. These findings highlight a previously underappreciated mechanism by which cancer cells exploit their mechanical environment to drive disease progression and offer new targets for therapeutic intervention.

## Acknowledgements

The authors greatly acknowledge the Cell and Tissue Imaging (PICT-IBiSA), Institut Curie, member of the French National Research Infrastructure France-BioImaging (ANR10-INBS-04). This work was supported by institutional grants from the Curie Institute, Institut National de la Santé et de la Recherche Médicale (INSERM), Centre National de la Recherche Scientifique (CNRS), and by specific grants from the Fondation pour la Recherche Médicale (FRM N° DGE20111123020), the Cancerople- IdF (n°2012-2-EML-04-IC-1), InCA (Cancer National Institute, n° 2011-1-LABEL-IC-4, PLBIO21-176, PLBIO-116), Fondation ARC pour la Recherche sur le Cancer to C.L and J.G. (Progamme Labellisé PGA1 RF20170205456), Agence Nationale de la Recherche (ANR-20-CE13-0002-01, ANR-19-CE15-0020-02 and ANR-24-CE44-4520-01), the CurieCoreTech Metabolomics and Lipidomics platform, the animal facility of CRBS, and by the European Union, EVCA Twining Project (Horizon GA n° 101079264). L.B. is supported by a Ph.D. fellowship from FRM. E.M. is the recipient of a Fondation ARC Passerelle grant. Work in the laboratory of J.G. is further supported by INSERM and University of Strasbourg, and by Fondation ARC to V.H. The authors would like to thank Dr. Clotilde Théry and Dr. Guillaume van Niel for the fruitful discussions and advice during the development of the present work. This work was performed as part of C.S’s doctoral thesis supported by grants from Fonds France Canada pour la Recherche (FFCR), Fondation ARC pour la Recherche sur le Cancer (Grant # ARCDOC42022010004536), and Becas Chile Doctorado en el Extranjero #72210467.

## Materials and Methods

### Cell lines and mice

HeLa, Hs578t, MDA-MB-231 cells along with the Cav1^-/-^ and cavin1^-/-^ sub lines were cultured in Dulbecco’s modified Eagle’s medium (DMEM-Glutamax, Gibco), with 10% FBS (FBS, Gibco), and 100U/ml penicillin and 100 μg/ml streptomycin (Gibco). Cell lines were grown at 37°C, under 5% CO2. Cell spheroids were prepared following the classical agarose cushion protocol. First, 50 µl of agarose 1,5% w/v in PBS (ultrapure agarose, Invitrogen) and 50 µl per well were dispensed in 96-well plate and incubated for 10-15 min for polymerization at room temperature. Then, cells were seeded on agarose cushion at 105 cells per well. Spheroid formation usually takes between 24 and 48 h. Cav1 and PTRF knockout cells were obtained by CRISPR-Cas9 gene editing.

Mice were 8-week-old BALB/c females that were housed and handled at CRBS (agreement number: #C67-482-35) according to the guidelines of INSERM and the ethical committee of Alsace (CREMEAS), following French and European Union animal welfare guidelines (Directive 2010/63/EU on the protection of animals used for scientific purposes).

All procedures were performed in accordance with French and European Union animal welfare guidelines and supervised by local ethics committee under the authorization number: APAFIS #38090.

### Antibodies and plasmids

Primary antibodies used include rabbit anti-cav1 (BD Transduction Laboratories, cat. no. 610059; 1:2,000 for WB and 1:200 for IF), mouse anti-Hrs (Abcam, cat. no. ab56468; 1:2,000 for WB), rabbit anti-STAM1/2 (Abcam, cat. no. ab76061; 1:2,500 for WB), mouse anti-CD9 (clone MM2/57; Sigma, cat. no. CBL162; 1:1,000 for WB), mouse anti-α-tubulin (clone B512; Sigma, cat. no. T5168; 1:5,000 for WB), rabbit anti-LC3B (Abcam, cat. no. ab51520; 1:3,000 for WB), mouse anti-CD63 (Santa Cruz, cat. no. sc-5275; 1:2,000 for WB and 1:100 for IF), mouse anti-calnexin (BD transduction laboratories. cat. no. 610523; 1:2,000 for WB), rabbit anti-PTRF (Proteintech. cat. no. 18892-1-ap. 1:1,000 for WB). rabbit anti-integrin αV (Cell signaling cat. no. 4711), rabbit anti-integrin α6 (Cell signaling cat. no. 3750), rabbit anti-integrin β5 (Cell signaling cat. no. 3629). Secondary antibodies used include mouse-HRP (Jackson ImmunoResearch, cat. no. 715-035-151; 1:5,000 for WB) and rabbit-HRP (Jackson ImmunoResearch, cat. no. 711-035-152; 1:5,000 for WB), mouse-Alexa 488 (Invitrogen, cat. no. A21202; 1:200 for IF), mouse-Cy3 (Jackson ImmunoResearch, cat. no. 715-166-150; 1:200 for IF), rabbit-Alexa 488 (Invitrogen, cat. no. A21206; 1:200 for IF), rabbit-Cy3 (Jackson ImmunoResearch, cat. no. 111-166-045; 1:200 for IF).

CAV1-K*R-mEGFP was a gift from Dr. Ari Helenius (Addgene plasmid # 27766; http://n2t.net/addgene:27766; RRID:Addgene_27766). The pCMV-CD63-pHluorin construct (Addgene, #130901) was transfected in HeLa cells using the lipofectamine LTX reagent (Invitrogen) protocol, with 1 µg of plasmid DNA. Cells were imaged 48 h after transfection.

### RNA interference

For HRS knockdown we used the following: Control siRNA (Dharmacon, Thermo Fisher, cat. no. SI03650325; 5′-AATTCTCCGAACGTGTCACGT-3′), the Hrs pool of four siRNA SMARTpool ON-TARGETplus HGS siRNA (Thermo Fisher, L-016835-00-0005; 5′-GAGGUAAACGUCCGUAACA-3′, 5′-GCACGUCUUUCCAGAAUUC-3′, 5′- AAAGAACUGUGGCCAGACA-3′ and 5′-GAACCCACACGUCGCCUUG-3′). Transfection was done using Lipofectamine RNAiMAX according to the manufacturer’s protocol (Thermo Fisher). siRNA was used at 20 nM. Depletion efficiency was assessed by immunoblotting.

### Mechanical stress models

To induce swelling of cells in 2D cultures we used hypo-osmotic shock. Hypo-osmotic shock was performed on cells by using the corresponding growth medium diluted appropriately in deionized water (1:9 dilution for 30 mOsm hypo-osmotic shock) for 5 minutes, after which media was changed to isosmotic media for EV collection.

To induce compression of multicellular spheroids, we used osmotic stress as described previously (Dolega et al., 2017). Shortly, we prepared hyperosmotic medium by adding high molecular weight dextran to reach a final concentration of 160 g/ml (2X solution). We used 2 MDa dextran (Sigma Aldrich, 95771) to avoid the penetration of the polymer in the cellular spheroid. Dextran containing medium was added on spheroid at 80 g/ml final concentration. For short-term compression experiments, spheroids were kept in dextran media for 5 min or 4 hours. For long-term compression experiments, spheroids were kept in dextran media for 1 or 5 days, and dextran containing culture medium is renewed after 3 days

### EV isolation

For the hypo-osmotic shock experiments, 6 × 10^6^ of cells were seeded per T75 culture flask. After 24 h, cells were washed 1 X with PBS (Gibco) and 12 ml of serum free iso or hypo-osmotic culture medium. The medium was changed after 5 min to an iso-osmotic, serum free culture medium, and cells were incubated for 1, 2, or 4 hours depending on the experiment, prior to EV isolation. For the osmotic-induced compression experiments, spheroids were washed with 1X PBS (Gibco) and 100 µl of serum free medium, with or without dextran for 5 minutes, 4 hours, 1 day, or 5 days depending on the experiment, prior to EV isolation. After treatments with mechanical stimulations (hypo-osmotic shock or osmotic induced compression) culture supernatant was collected and was subjected to serial centrifugations (2,000 g for 10 min, 11,000 g for 30 min at 4°C), followed by ultracentrifugation at 100,000 g for 90 min at 4°C (45Ti or TLA110 rotors, Beckman Coulter). The pellet containing EVs was washed in cold 1x PBS (Gibco) and ultracentrifuged again at 100,000 g for 90 min at 4°C.

### EV characterization by NTA

NTA was performed using ZetaView PMX-120 (Particle Metrix) equipped with a 488 nm laser, with software version 8.05.02 to measure the concentration and distribution size of particles by evaluating the Brownian motion in a light scattering system or the NanoAnalyzer—NanoFCM, S/N FNAN30E22101948 with single-photon counting modules to detect the side scatter and green fluorescence of individual particles. The samples were diluted in 1X PBS (Gibco) to obtain a concentration of 10^6^ to 10^9^ particles/ml per field.

### Western blotting

Cell and EV extracts were separated by SDS-PAGE, transferred to nitrocellulose, blocked in 1x PBS containing 3% BSA or 5% nonfat milk according to the antibody used. Membranes were probed overnight at 4°C. Bound antibodies were developed using SuperSignal™ West Femto Maximum Sensitivity Substrate (Thermo Scientific) and the ChemiDoc Touch Imaging System (BioRad). Protein bands were quantified by densitometric analysis using the ImageJ

### Electron microscopy

For imaging cell spheroids, epon embedding was used to preserve the integrity of cell structures for electron microscopy (EM). Spheroids were fixed sequentially for 1 h at room temperature with 1.25% glutaraldehyde in 0.1 M Cacodylate and then overnight at 4 °C. Cells were washed extensively with 0.1 M Cacodylate, pH 7.2. Post-fixation was performed for 1 h at room temperature with 1% OsO4 in 0.1 M Cacodylate, pH 7.2. Spheroids were dehydrated through a graded-concentration series of ethanol (50, 70, 90, then 100%, each for 10 min at RT). Embedding was finally performed in LX112 resin. Cells were infiltrated with an increasing ratio of LX112: ethanol solution (1:2, 1:1 and 2:1) and finally with pure LX112. Samples in resin were polymerized overnight at 60°C. Semi-thin 500 nm sections were sliced using a Leica UCT ultra microtome and mounted onto microscopic glass slides and dried on a hot plate. Semithin sections were stained for 30 s on a hot plate with a mix of Azure B and basic fuchsin in sodium tetraborate (Morikawa et al., 2018). Sections were then mounted in DPX for microscopy and covered with coverslips. Micrographs were acquired on an Upright Wide field Leica DM6000b Microscope equipped with a color CoolSNAP HQ2 camera. Ultrathin 65 nm sections were sliced using a Leica UCT ultra microtome and mounted on nickel formvar/carbon-coated grids for observations. Contrast was obtained by incubation of the sections for 10 min in 4% uranyl acetate followed by 1 min in lead citrate. Caveolae were identified based on their ultrastructural features. The length of plasma membranes observed were measured using ImageJ software and the number of the structures observed was reported to µm of membrane.

EV imaging was performed as previously described (Corona et al., 2023). Briefly, EVs were deposited on a coated-side of a formvar-carbon grid and left to adhere. Grids were incubated in uranyl acetate on ice and in the dark. Grids were removed from the droplet then air dried for 30 mins on the loops. Electron micrographs were acquired on a Tecnai Spirit electron microscope (FEI, Eindhoven, The Netherlands) equipped with a 4k CCD camera (EMSIS GmbH, Münster, Germany).

### Immunofluorescence

After the treatments, cells were fixed with 4% paraformaldehyde (PFA) (EMS) for 15 min at room temperature. Cells were incubated for 1h in a blocking solution: 1x PBS with 0.1% triton and 0.3% BSA. Then primary and secondary antibodies were successively incubated during 1 h each at RT in 1x PBS containing 0.1% triton and 0.1% BSA. Coverslips were then mounted on microscope slides with Fluoromount G (Invitrogen). Images were acquired on a Zeiss LSM 780 confocal microscope with a 63x/1.46 Oil objective. At least 10 cells per replicate were imaged.

### Live TIRF microscopy

Cells were grown in 35-mm imaging plates (FluoroDish WPI; IBIDI). An inverted Eclipse Ti-E (Nikon) full motorized with PFS 2 (Perfect Focus System) to reduce the drift was used with a 100 ×CFI PlanApoTIRF, oil, 1.49/0.12-mm objective (Nikon), images were acquired with Metamorph (Molecular Devices). A cage incubator (Life Imaging Services) was used to keep the imaging chamber at 37°C and 5% CO2. To visualize fusion events, 1-3 min videos were acquired in different areas to analyze at least 20 cells per condition.

### Sample preparation for lipidomics

After EV isolation by ultracentrifugation, the EV pellet was reconstituted in 200 µl of 150 mM ammonium bicarbonate (NH4HCO3), then quickly frozen with liquid nitrogen and stored at -80 °C or later use. Lipids were extracted in chloroform and methanol pre-mixed with internal standard mix, then analysed at the CurieCoreTech Metabolomics and Lipidomics platform.

### EV collection for MS-based proteomics

2,5x10^6^ cells per 10 cm dish were plated the day before the osmotic shock. For each osmotic experiment, medium was collected from 10 plates, while for resting cells, medium from 15 plates. The medium was centrifuged at 2,000 g for 15 min, and EVs were purified by ultracentrifugation, following established protocols. The concentration of recovered EVs was measured by nanoparticle tracking analysis (ZetaView, Particle Metrix) and resuspended to a final concentration of 1x10^11^ particles in 10 µl of PBS. Five independent biological replicates were analyzed by liquid chromatography Tandem mass spectrometry (LC-MS/MS).

#### Sample Preparation for LC-MS/MS

Protein pellets of 10 µg each were dried under vacuum using a Savant Centrifuge SpeedVac concentrator (Thermo Fisher Scientific). The dried pellets were then solubilized and reduced in 10 µL of 8 M urea, 100 mM ammonium bicarbonate, and 5 mM dithiothreitol (DTT) at pH 8.0, and incubated at 57°C for 30 min. Following this, iodoacetamide was added to a final concentration of 10 mM. The alkylation reaction was conducted in the dark for 30 minutes at room temperature. The samples were subsequently diluted to a final urea concentration of 1 M using 100 mM ammonium bicarbonate at pH 8.0. Protein digestion was performed using trypsin/LysC (0.2 µg, Promega) in a total volume of 100 µL, with the mixture incubated overnight at 37°C under constant vortexing. To desalt the digested peptides, the samples were passed through homemade C18 StageTips. Elution of peptides was achieved using a solution of 40% acetonitrile (CH3CN) and 60% water containing 0.1% formic acid. The eluate was then concentrated to dryness under vacuum.

#### LC-MS/MS analysis

LC was performed with a RSLCnano system (Ultimate 3000, Thermo Scientific) coupled to an Orbitrap Exploris 480 MS, interfaced by a Nanospray Flex ion source (Thermo Scientific). Peptides were trapped on a C18 column (75 μm inner diameter × 2 cm; nanoViper Acclaim PepMap^TM^ 100, Thermo Scientific) with buffer A (2/98 CH3CN/H2O in 0.1% formic acid) at a flow rate of 2.5 µL/min over 4 min. Separation was performed on a 50 cm x 75 μm C18 column (nanoViper Acclaim PepMap^TM^ RSLC, 2 μm, 100Å, Thermo Scientific) regulated to a temperature of 50°C, and separated with a linear gradient from 98% buffer A (100% H2O + 0,1% formic acid) to 30% buffer B (100% CH3CN + 0,1% formic acid) at a flow rate of 300 nL/min over 91 min. The mass spectrometer was operated in data-independent analysis (DIA). MS full scans were performed in the Orbitrap mass analyzer in mass range 375-1500 m/z with a resolution of 120,000 at m/z 200, a normalized AGC target set at 300% and a maximum injection time set at custom. 40 DIA scans of 15 Da isolation window from 400 to 1000 m/z are generated with a resolution of 15000, NCE at 25, normalized AGC target at 3000% and a maximum injection time set at Auto.

#### Data processing

For identification, the data were searched against the Homo sapiens (UP000005640) Uniprot database using Spectronaut v17 (Biognosys) by directDIA+ analysis using default search settings. Enzyme specificity was set to trypsin and a maximum of two missed cleavage sites was allowed. Carbamidomethyl, N-terminal acetylation and oxidation of methionine were set as variable modifications. The resulting files were further processed using myProMS v3.10. https://github.com/bioinfo-pf-curie/myproms (Poullet et al., 2007).

For protein quantification, ion XICs (sum of fragment peak areas) from proteotypic peptides shared between compared conditions (TopN matching) were used, with missed cleavages and no modification allowed. Median and scale normalization at peptide level was applied on the total signal to correct the XICs for each biological replicate (N = 5). To evaluate the statistical significance of the change in protein abundance, a linear model (adjusted on peptides and biological replicates) was performed, and a two-sided *t*-test was applied on the fold change estimated by the model. The p-values were then adjusted using the Benjamini–Hochberg FDR procedure. Proteins with at least two distinct peptides in three replicates of a same state, a 1.5-fold enrichment and an adjusted p-value ≤ 0.05 were considered significantly enriched in sample comparisons. Proteins unique to a condition were also considered if they matched the peptides criteria. Proteins selected with these criteria were further used for Gene Ontology (GO) enrichment analysis using all quantified proteins as background set. LFQ was performed following the algorithm as described (Cox et al., 2014), with the minimum number of peptide ratios set to 2 and with large ratios stabilization feature used. Protein LFQ values were further normalized (median and scale) to correct for potential remaining total intensity biases.

The mass spectrometry proteomics raw data have been deposited to the ProteomeXchange Consortium via the PRIDE (Perez-Riverol et al., 2025) partner repository with the dataset identifier PXD065554 (, password:1uZ6TUGHjnoB)

### Migration and invasion assays

Migration assays of Hs578t cells were performed by seeding 300,000 cells in each well of a 24-well plate 24 h before the experiment. The cells were incubated with 1 μg/ml of EVs 37 °C for 4 h. After incubation, a scratch was done in the middle of each well with a pipette tip, cells were washed 1 time with PBS and incubated with FBS-free culture medium. Cell migration was monitored using time-lapse microscopy (IncuCyte Live Cell Analysis Systems, 4x objective lens, Sartorius) with an interval of 3 h during 21 h. Image analysis was performed with ImageJ using a wound healing size tool (Suarez-Arnedo et al., 2020).

Invasion assays were performed in 8 μm pore size 24-well inserts (Greiner bio-one) pretreated with Matrigel (BD Biosciences) according to the manufacturer’s instructions. Briefly, the inserts were coated with Matrigel and were left in a 37°C incubator for 30 mins before seeding. Then, 300,000 Hs578t cells previously treated with 1 μg/ml of EVs during 24 h were resuspended in serum-free medium and added to the top of each chamber insert, and medium with FBS as a chemo-attractant was added to the bottom chamber. After 24 h, inserts were removed, washed and cells that migrated to the lower side of the inserts were stained with 0.1% toluidine blue, washed, and counted in an inverted microscope.

### EV staining and uptake

EVs isolated by ultracentrifugation were labeled by incubation with 20 µM CFDA-SE (Thermofisher) for 2 h at 37°C. To eliminate surplus dye, size-exclusion chromatography (SEC) was performed using Exo-spin mini columns (Interchim). CFSE-labeled EVs were then collected, and their particle concentration and fluorescence intensity were analyzed using Nanoparticle Tracking Analysis (NTA) and Nano Flow Cytometry (NanoFCM).

For the uptake experiments, acceptor cells were seeded 24 h before at a density of 100,000 cells per well in a 24-well plate on coverslips. The acceptor cells were then incubated with CFSE-labeled EVs at a final concentration of 5x10^9^ EVs/mL for 4 h at 37 °C. After incubation, the cells were washed with PBS and fixed with PBS 4% paraformaldehyde in PBS for 15 mins at room temperature. The cells were then permeabilized for 30 mins at room temperature in PBS containing 1% bovine serum albumin (BSA) 0.1% Triton-X-100. Cells were stained with WGA-rhodamine (ThermoFischer) for membrane visualization. Image analysis and colocalization quantification were performed using ImageJ software (NIH, Maryland, USA).

For the organotropism experiments EV staining was performed using MemGlow-Cy5 dye (500nM), as previously described, for 20 min at room temperature in the dark (with up-downs to mix everything every 4 min), and washed in PBS in an Amicon 10kDa for 20 min at 4000g (Hyenne et al., 2019).

### *In vivo* organotropism

Mice were injected with 3x10^9^ stained EVs in the lateral tail vein and were euthanized 45 min post-injection by cervical dislocation. All the organs were dissected and placed in a holder for imaging with the In Vivo Imaging System (IVIS, Perkin) using the fluorescent mode with 20 seconds excitation, binning 1, F/Stop 1,2 with the Cy5 channel, images were analyzed using the Living Image Software (Perkin).

### Statistical analysis

All analyses were performed using GraphPad Prism version 6.0 to 8.0, (GraphPad Software). Two-tailed (paired or unpaired) Student’s t-tests were used when comparing only two conditions. For more than two conditions, the Kruskal–Wallis test, ordinary one-way ANOVA or two-way ANOVA were used. Significantly different comparisons of means are marked on the graphs with asterisks. Error bars denote SEM.

---

**Supplementary figure 1:**
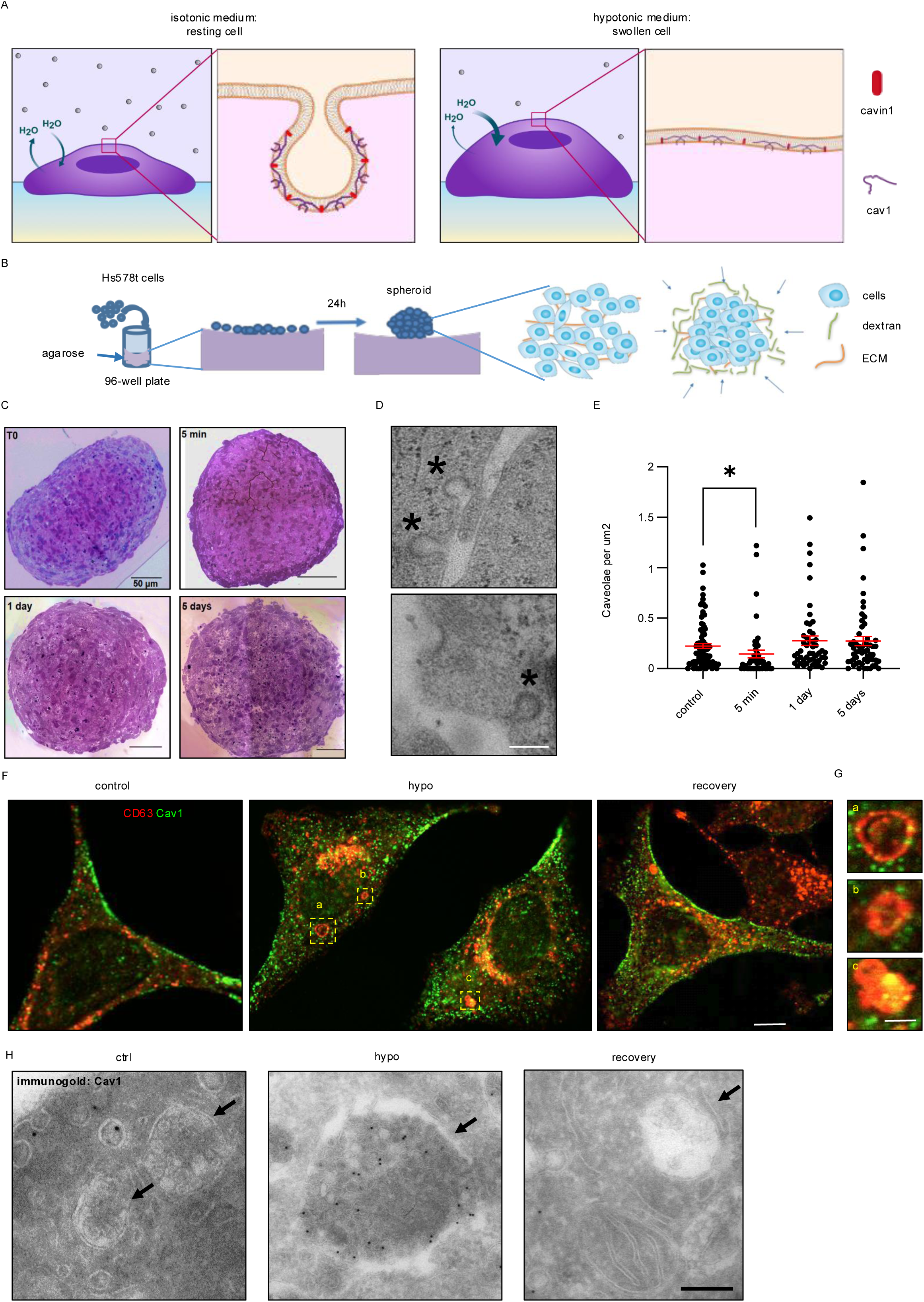
Validation of mechanical stress models and their impact on caveolae and multivesicular bodies. **A.** Schematic representation of the effect of hypo-osmotic shock on caveolae. In resting conditions, caveolae are fully budded invaginations where Cav1 and cavin1 can interact. Upon hypo-osmotic shock, caveolae flatten out, resulting in the diffusion of non-caveolar Cav1 at the plasma membrane and loss of interaction between Cav1 and Cavin1. This process is reverted upon return to resting conditions **B.** Schematic representation of the dextran compression model on spheroids. Cells are seeded on an agarose bed and left to spontaneously form spheroids. High molecular weight dextran is used to induce osmotic induced mechanical compression of the spheroids. **C.** Representative images of Hs578t spheroids at resting conditions and after 5 min, 1 day and 5 days of mechanical compression. **D.** Representative EM images of caveolae structures (*) observed on spheroids. **E.** Quantification of caveolar structures in Hs578t spheroids at resting conditions and after 5 min, 1 day and 5 days of mechanical compression. Graph represents the data in caveolae per area. **F.** Representative immunofluorescence of Cav1 and CD63 in HeLa cells in resting conditions, 5 min after exposure to hypo-osmotic shock, and 10 minutes after recovery in iso-osmotic media. Scale bar 5 μm. **G.** Magnified inserts from B showing enlarged endosome-like structures in the cell after osmotic shock. Scale bar 1 μm. **H.** Representative EM images of MVBs in MLEC cell in resting conditions, during hypo-osmotic shock and in recovery. Immunogold labeling against Cav1. Arrows indicate MVB structures. Scale bar 200 nm. * *P* < 0.05.

**Supplementary figure 2:**
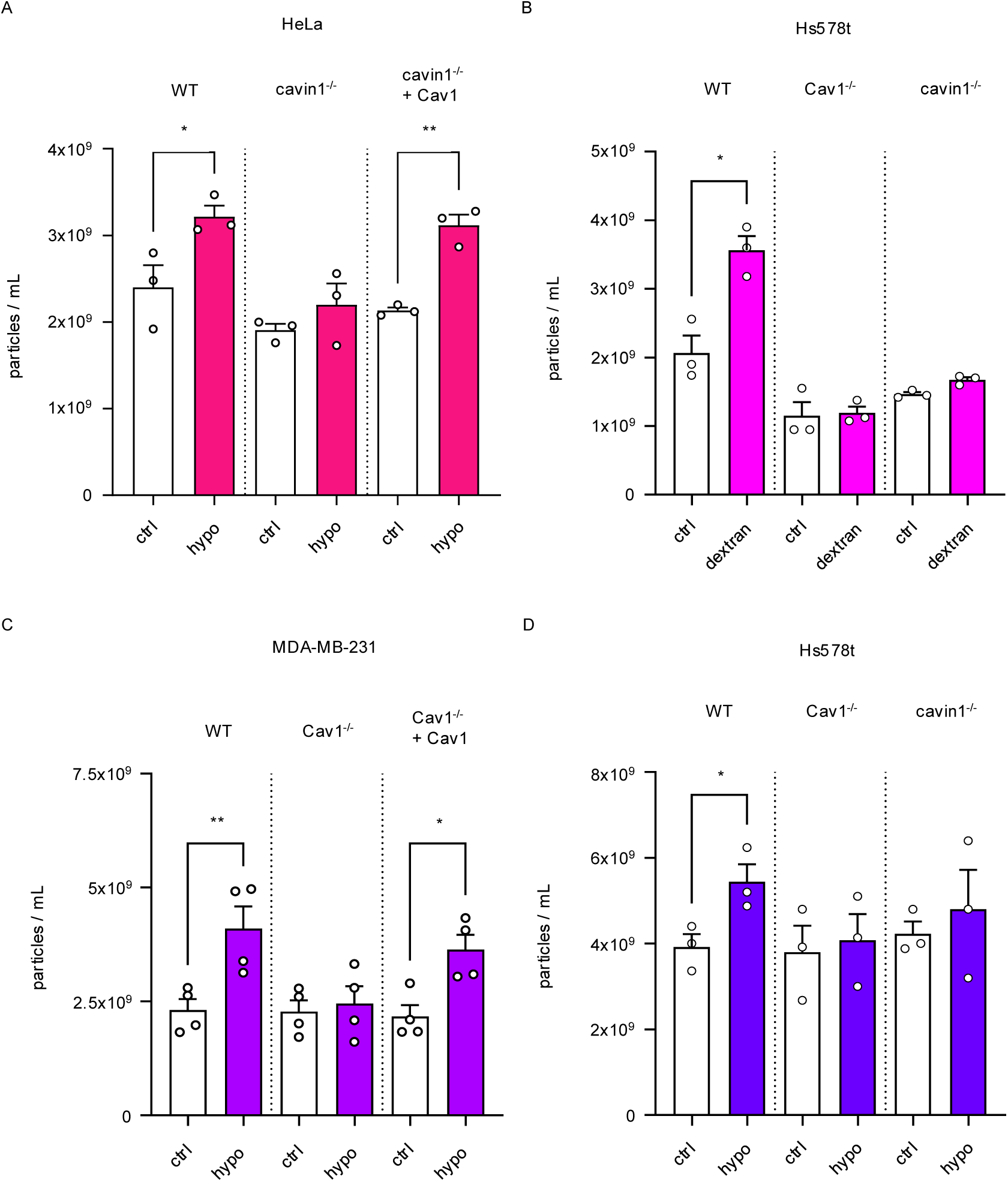
Quantification of EV release following mechanical stress in diverse cell lines with modulated caveolae constituent expression. **A.** Quantification of EVs released by WT, cavin1^-/-^, and cavin1^-/-^ overexpressing Cav1 HeLa cells after exposure to hypo-osmotic shock. Graph shows total secreted EVs. **B.** Quantification of EVs released by WT, Cav1^-/-^ and cavin1^-/-^ Hs578t cells after dextran compression. Graph shows total secreted EVs. **C.** Quantification of EVs released by WT, Cav1^-/-^ and Cav1^-/-^ with re-expression of Cav1 MDA-MB-231 cells after exposure to hypo-osmotic shock. Graph shows total secreted EVs. **D.** Quantification of EVs released by WT, Cav1^-/-^ and cavin1^-/-^ Hs578t cells after exposure to hypo-osmotic shock. Graph shows total secreted EVs. ** *P* < 0.01, * *P* < 0.05.

**Supplementary figure 3:**
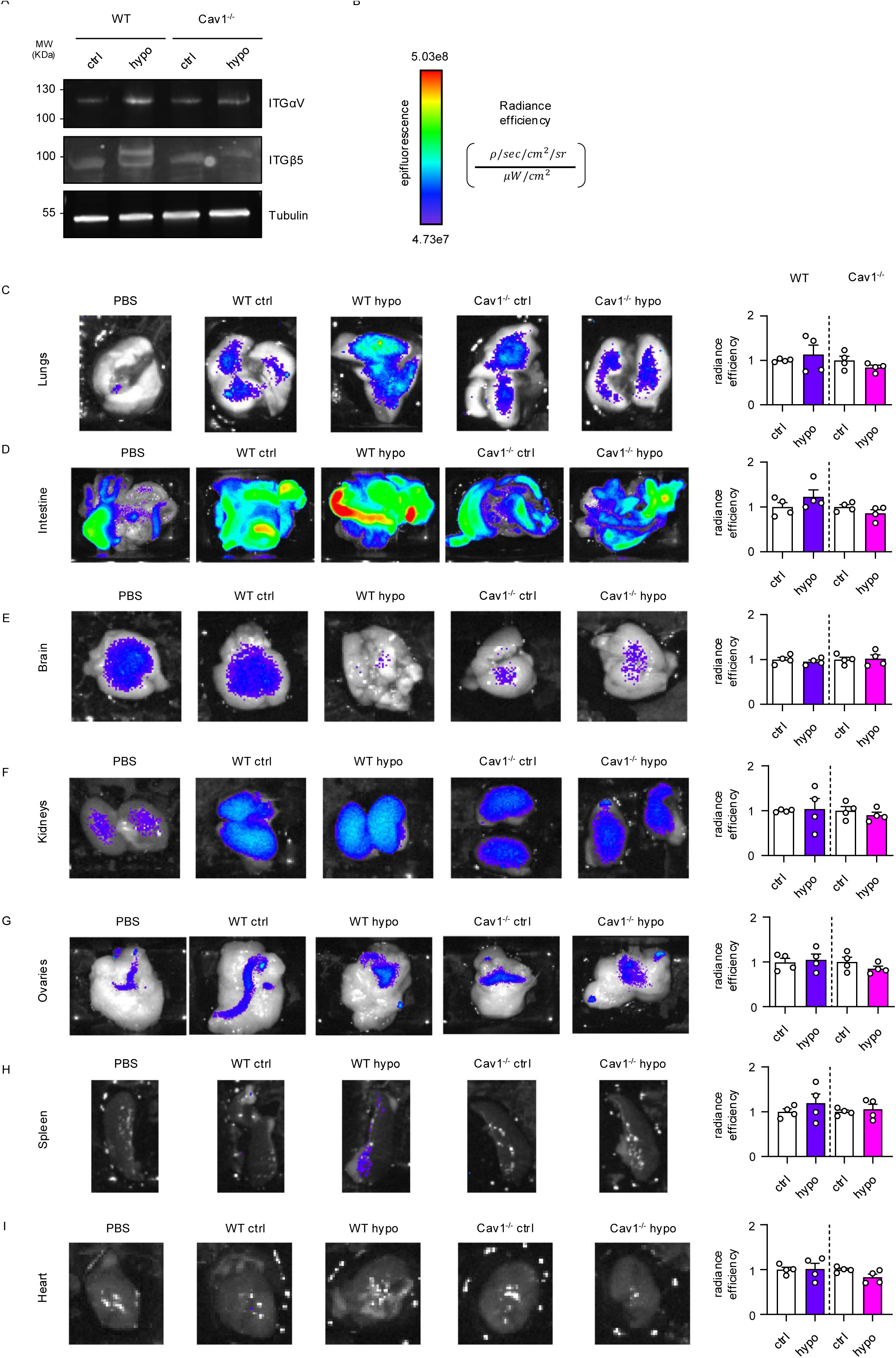
Integrin expression and organ-specific biodistribution of EVs under mechanical stress. **A.** Western blot analysis of integrins αV and β5 in EVs from WT and Cav1^-/-^ EVs. **B.** Formula used to calculate radiance efficiency. **C-I.** Representative epifluorescence images showing EV accumulation in various organs: lungs (C), intestine (D), brain (E), kidneys (F), ovaries (G), spleen (H), and heart (I). Quantification of radiance efficiency is shown in the graph on the right.

**Supplementary figure 4:**
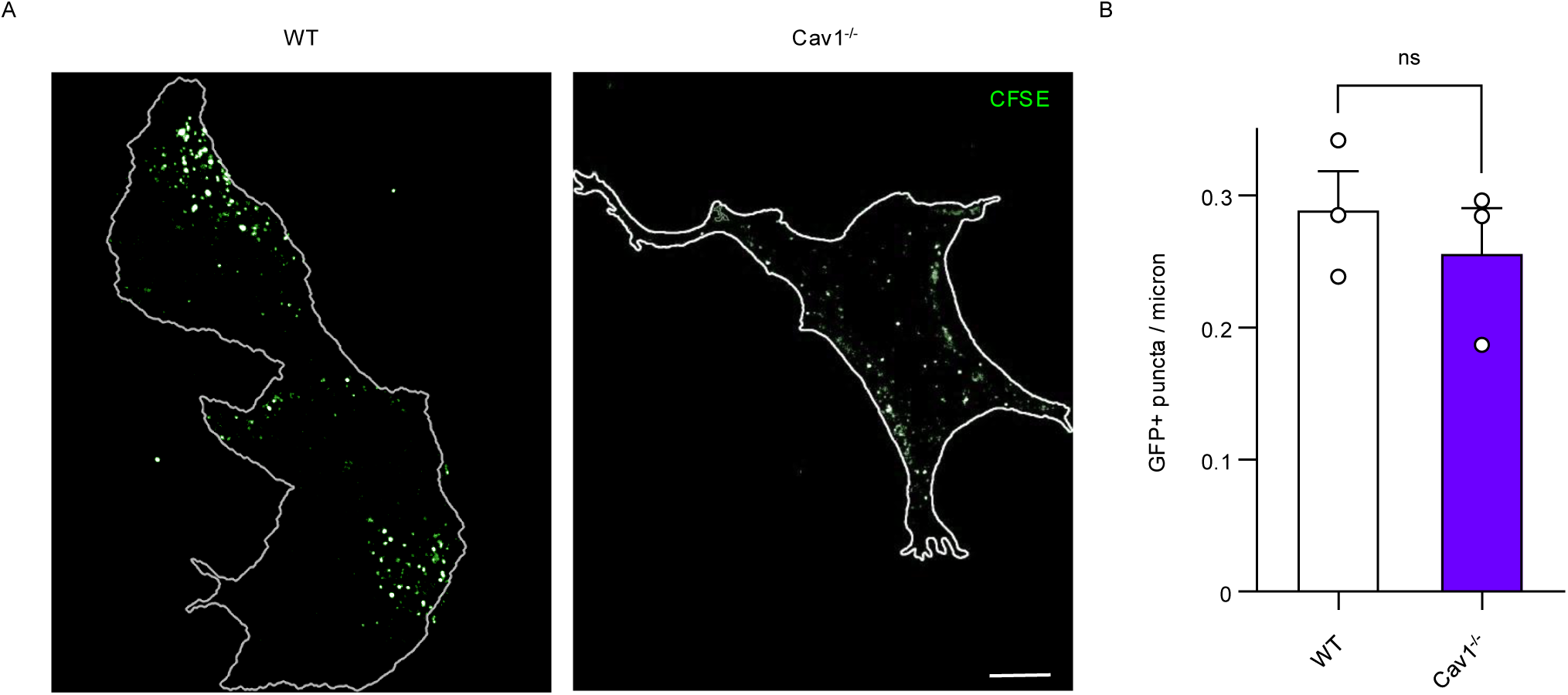
Equal internalization of WT and Cav1-deficient EVs by Hs578t cells. **A.** Representative confocal images of Hs578t acceptor cells after a 4 h incubation with CFSE-labeled EVs. Scale bar 5 μm. **B.** Quantification of CFSE-positive dots per cell in WGA-labeled Hs578t WT cells after a 4 h incubation with CFSE-EVs from WT or Cav1-depleted HeLa cells. Each dot represents the number of CFSE-positive dots per micron.

## Notes

### Competing Interest Statement

The authors have declared no competing interest.

### Summary of Updates

This version of the manuscript includes new experimental results that strengthen the main conclusions. Most figures have been updated with new data or reordered to improve clarity. Figure 4 has been substantially revised to incorporate new findings, and Figure 7 now includes additional results related to organotropism. Supplementary Figure 3 has also been updated to show additional organs analyzed in the organotropism experiments. Additionally, new contributing authors have been added to reflect their involvement in the revised work.

